# Sustainable land management enhances ecological and economic multifunctionality under ambient and future climate

**DOI:** 10.1101/2023.01.30.525902

**Authors:** Friedrich Scherzinger, Martin Schädler, Thomas Reitz, Rui Yin, Harald Auge, Ines Merbach, Christiane Roscher, Stan Harpole, Sigrid Berger, Evgenia Blagodatskaya, Julia Siebert, Marcel Ciobanu, Nico Eisenhauer, Martin Quaas

## Abstract

Anthropogenic activity is threatening ecosystem multifunctionality, i.e. the ability of ecosystems to provide multiple functions and services which are vital for human well-being. Here we assess how multifunctionality of agroecosystems in Central Germany depends on land-use type and climate change. Our analysis included 13 ecosystem functions in a large-scale field experiment with five different land-use types (three grassland and two farmland types either sustainably or intensively managed) under two different climate scenarios (ambient and future climate). We consider ecological measures of multifunctionality using averaging approaches with different weights, i.a. reflecting preferences of farmers and environmentalists, and assess an economic multifunctionality measure based on the total value of ecosystem services. Results show that intensive management and future climate decrease ecological multifunctionality for multiple weighting scenarios in both grassland and farmland. Only under a weighting according to farmers’ preferences, intensively-managed grassland shows higher multifunctionality as compared to sustainably-managed grassland. The economic multifunctionality measure, which includes economic benefits for society at large, reveals a multifunctionality about ∼1.7 times higher for sustainable compared to intensive management for both grassland and farmland. Above-belowground biodiversity correlates positively with ecosystem multifunctionality and is expected to be one of its main drivers. Based on these findings, we suggest to promote and economically incentivise sustainable land management that enhances both ecological and economic multifunctionality, also under future climatic conditions.

## 2. Introduction

Ecosystem functions are natural processes (biological, geochemical, and physical) that are indirectly linked to the provision of ecosystem services and the economic value they generate for ecosystem managers and society at large (Groot et al., 2002). Ecosystem multifunctionality (EMF) describes the ability of ecosystems to provide multiple functions or services simultaneously (Manning et al., 2018). EMF research puts into focus the trade-offs and synergies that occur between different ecosystem functions (Dooley et al., 2015). Due to the interdependent and overlapping nature of different services and functions, EMF is more than just the sum of its functions and services, and a loss in one EMF component cannot necessarily be fully compensated by an increase in another (Manning, 2017). How to measure ecological multifunctionality depends on stakeholders’ preferences for different functions and associated ecosystem services (Allan et al., 2015). Here, we contrast alternative ecological approaches of measuring EMF, and an economic approach that is based on the total economic value of multiple ecosystem services driven by EMF.

EMF is affected by land use as well as by anthropogenic climate change. Climate change is expected to show spatially diverse, but net negative effects on ecosystem services (Scholes, 2016; IPCC, 2014). In the context of agriculture, biodiversity loss due to land-use intensification (increased application of agrochemicals and machinery) is a major driver of a decrease in EMF (Allan et al., 2015; Hector & Bagchi 2007). Here, the role of biodiversity is ambiguous as it is both a driver of many different ecosystem functions and a function (or service) by itself. Biodiversity stabilises and increases many different natural processes, including ecosystem functions, and ecosystem services associated with them (Hong et al., 2022; Isbell et al., 2015).

Within the field of EMF research, most studies have focused on single drivers of multifunctionality (e.g., the effect of microbial diversity on (soil) multifunctionality (Delgado-Baquerizo et al., 2017; Wagg et al., 2014), the effect of biodiversity on multifunctionality (Pan et al., 2017; Soliveres et al., 2016), or the change of (soil) multifunctionality across a natural climate gradient (Ding & Eldridge, 2021).

Whereas overall effects of future climate on EMF have found little attention so far (Giling et al., 2019), different studies focus on the effect of biodiversity on EMF under different environmental conditions. Higher biodiversity was shown to increase EMF (Hong et al., 2022; Eisenhauer et al., 2018) while it remains unclear how the biodiversity-EMF relationship changes under future compared to ambient environmental conditions. No significant difference in the magnitude of the effect of biodiversity on soil EMF was shown under ambient compared to future climate conditions, whereas future climate was simulated with elevated CO_2_ concentration and enhanced nitrogen deposition (Eisenhauer et al., 2018). Contrary, a stronger effect of biodiversity on EMF was observed under more stressful environments produced by global change drivers (indicating future climate conditions) implying ecosystems with higher biodiversity being more resistant to future climate (Hong et al., 2022).

Some studies have focused on the overall effect of management intensity of agricultural grasslands on EMF (Neyret et al., 2021; Allan et al., 2015; Fischer et al., 2010). In farmlands, a higher EMF was shown for organic / conservation agricultural ecosystems (promoting regulating and supporting services) compared to conventional agriculture ecosystems (promoting a small number of provision services; Wittwer et al., 2021). Moreover, production with organic or conservation agriculture resulted in a higher economic value due to higher product prices (subsidies not included) (Wittwer et al., 2021). Despite the field of EMF research gaining more and more attention (Giling et al., 2019), the impact of different land-use types and future climate on EMF in the context of agriculture with a direct quantification of those changes in monetary units has not been assessed before. This study aims to address this research gap.

Therefore, we analysed data from the Global Change Experimental Facility (GCEF) in Saxony-Anhalt, Germany, a large field experiment with orthogonal manipulation of climate (ambient and future climate type) and land use (five different land-use types: three grassland and two farmland types either sustainably managed (no application of mineral nitrogen fertiliser and pesticides) or intensively managed (application of mineral nitrogen fertiliser and pesticides), see Table 1) within 50 plots of approximately 400 m^2^ each (Schädler et al., 2019).

**Table 1.** Overview of the land-use types considered in this study. Classification into sustainable and intensive management was done based on the input of agrochemicals (mineral nitrogen fertiliser and pesticides) which was refrained under sustainable management.

|  | **Sustainable management** | **Intensive management** |
| --- | --- | --- |
| Grassland | Extensive meadow (EM) | Intensive meadow (IM) |
|  | Extensive pasture (EP) |  |
| Farmland | Organic farming (OF) | Conventional farming (CF) |

We measured the levels of 13 ecosystem functions approximating five ecosystem services over seven years after establishment of the treatments (see Figure 1). The classification and categorization of ecosystem functions was done based on the Millennium Ecosystem Assessment Program (Millennium Ecosystem Assessment, 2005). Yield indicates **food production**; total organic soil carbon (yearly flux) indicates **climate regulation**; (absence of) nitrogen surplus indicates **water pollution control**; microbial biomass, enzymatic activity, and decomposition rate indicate **soil health preservation** as they enhance the nutrient release rate and the soil water retention (Dornbush et al., 2002), see materials and methods for details. Biodiversity of soil nematodes and meso- / macrofauna indicates **biodiversity conservation** that provides multidimensional values to people (Dasgupta, 2021). An increase in the level of the ecosystem functions is desirable (except for nitrogen surplus), as it indicates an increase in the respective ecosystem services.

**Figure 1.**
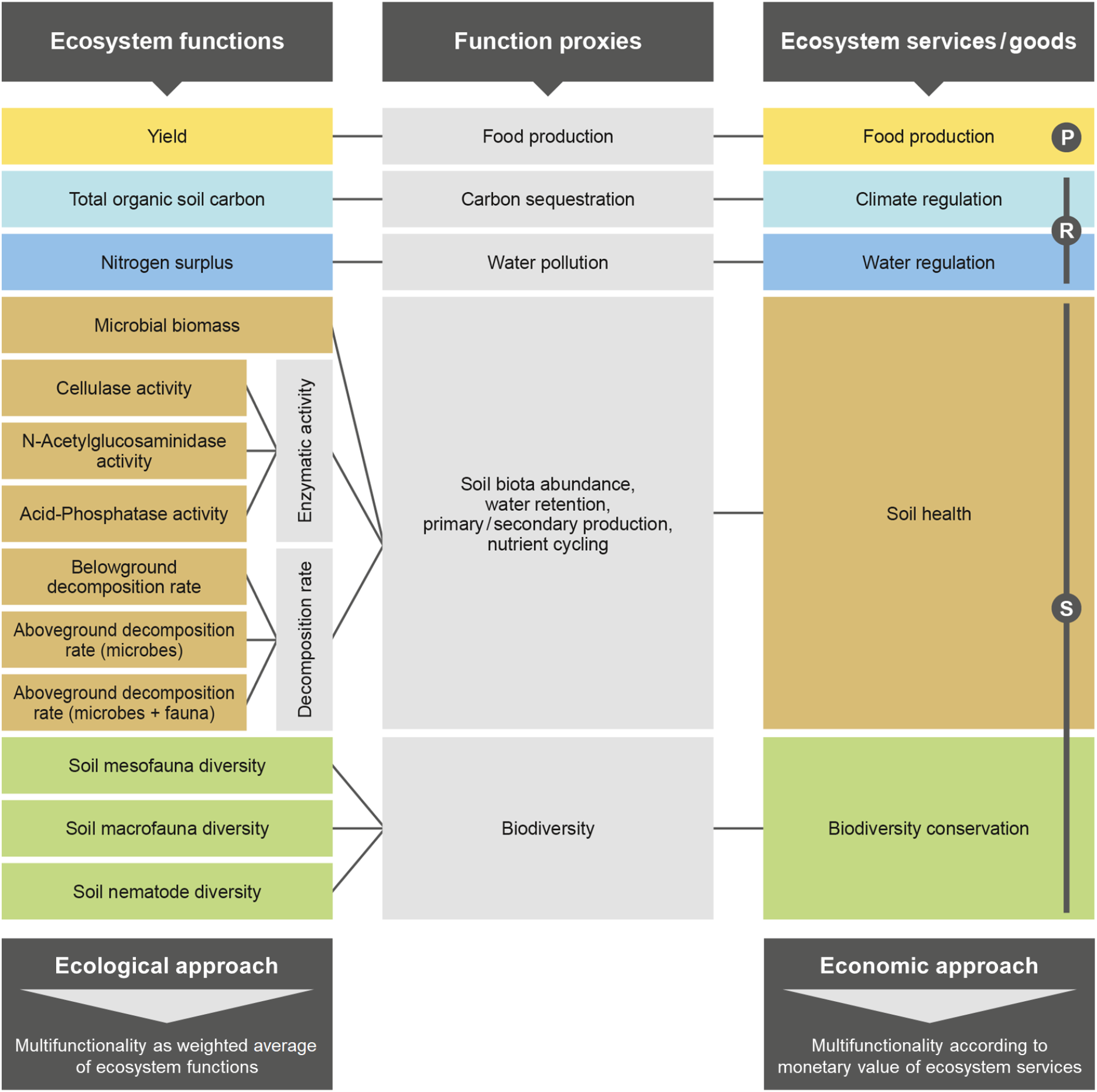
Conceptual framework for the calculation of ecosystem multifunctionality (EMF) based on 13 ecosystem functions approximating five ecosystem goods / services (P: provisioning service, R: regulating services, S: supporting services) with both an ecological and an economic approach.

To measure ecological EMF we used the averaging approach (Byrnes et al., 2014). We normalised all ecosystem function levels to values ranging between 0 and 1 based on the minimum and maximum observed values of the respective function within the experiment, and then calculated EMF on the plot level using the weighted average of the 13 ecosystem functions’ normalised levels. The weighting was done based on different objectives and stakeholder preferences (see Figure 7, Supplementary material). Main results exclude the first two years of the experiment, as many soil functions require time to respond to experimental treatments. For the calculation of EMF, the mean values of ecosystem function levels of the last five years of the experiment were used. For the economic approach of quantifying the value of EMF, we used the total economic value of the ecosystem services listed in Figure 1. For food production, climate regulation and water regulation, we computed the value by multiplying the quantity of the services with accounting prices from the literature. For biodiversity, we developed a model of quantifying the insurance value, which is an important part of the economic value of biodiversity (Farnsworth et al., 2015) due to its stabilising effect on direct use values (e.g., yield) (Augeraud-Véron et al., 2019; Baumgärtner & Quaas, 2010; Baumgärtner, 2007). A yield-stabilising effect was also described for soil health (Gao et al., 2015; Zhang et al., 2009; Watts & Torbert, 2011), leading to an insurance value that we quantified with a similar model.

Further, we tested if certain ecosystem functions drive EMF. To analyse biodiversity-EMF relationships, we assessed EMF based on the multiple threshold approach (Byrnes et al., 2014). The results of this study may provide decision-making support to implement strategies to control and counteract the effects of land-use intensification and climate change on EMF and to steer policy decisions accordingly.

## 3. Results

### 3.1. Effects of climate and land-use type on ecological multifunctionality measures

Ecosystem functions show different responses to the climate and land-use treatments (Section 8.1, Table 4, Figure 6, Supplementary material). The effect of the different climate and land-use types on ecosystem multifunctionality (EMF) is strongly dependent on the weighting scenario (Table 2, Figure 2). EMF is decreased under future climate for two out of four weighting scenarios (equal ecosystem function weighting and weighting according to farmers’ preferences). The land-use type has a strong effect on EMF under all weighting scenarios. Under equal weighting of all ecosystem functions, EMF of grassland is reduced under intensive management, whereas for farmland (where EMF is generally lower) no difference was found between the management types. Under equal weighting of all ecosystem services, for both grassland and farmland, intensive management decreases EMF. Under a weighting according to preferences of farmers, EMF is smaller for grasslands compared to farmlands, and for grassland, the intensive management scenario shows a higher EMF, whereas for farmland, EMF is higher under sustainable management. Under a weighting according to preferences of environmentalists, the sustainably-managed grassland types have significantly higher EMF than the other land-use types.

**Table 2.** General linear regression model table of F and p values (in brackets: numerator and denominator d.f.) of the effect of the two factors land-use type (LUT), climate type (Climate), and their interaction (LUT:Climate) on ecological ecosystem multifunctionality EMF for different weighting scenarios. Bold values indicate a significant effect of the respective factor or interaction (*** p<0.001; ** p<0.01; * p<0.05).

|  | LUT |  |  | Climate |  |  | LUT:Climate |  |
| --- | --- | --- | --- | --- | --- | --- | --- | --- |
| Weighting | F(4,32) | p value |  | F(1,8) | p value |  | F(4,32)<br>value | p value |
| Equal ecosystem function weighting | **44.61** | **0.000** | *** | **7.96** | **0.022** | * | 0.26 | 0.904 |
| Equal ecosystem service weighting | **23.11** | **0.000** | *** | 2.34 | 0.164 |  | 0.20 | 0.940 |
| Weighting according to farmers' preferences | **82.64** | **0.000** | *** | **8.17** | **0.021** | * | 0.603 | 0.700 |
| Weighting according to environmentalists' preferences | **75.24** | **0.000** | *** | 0.28 | 0.613 |  | 0.05 | 1.000 |

**Figure 2.**
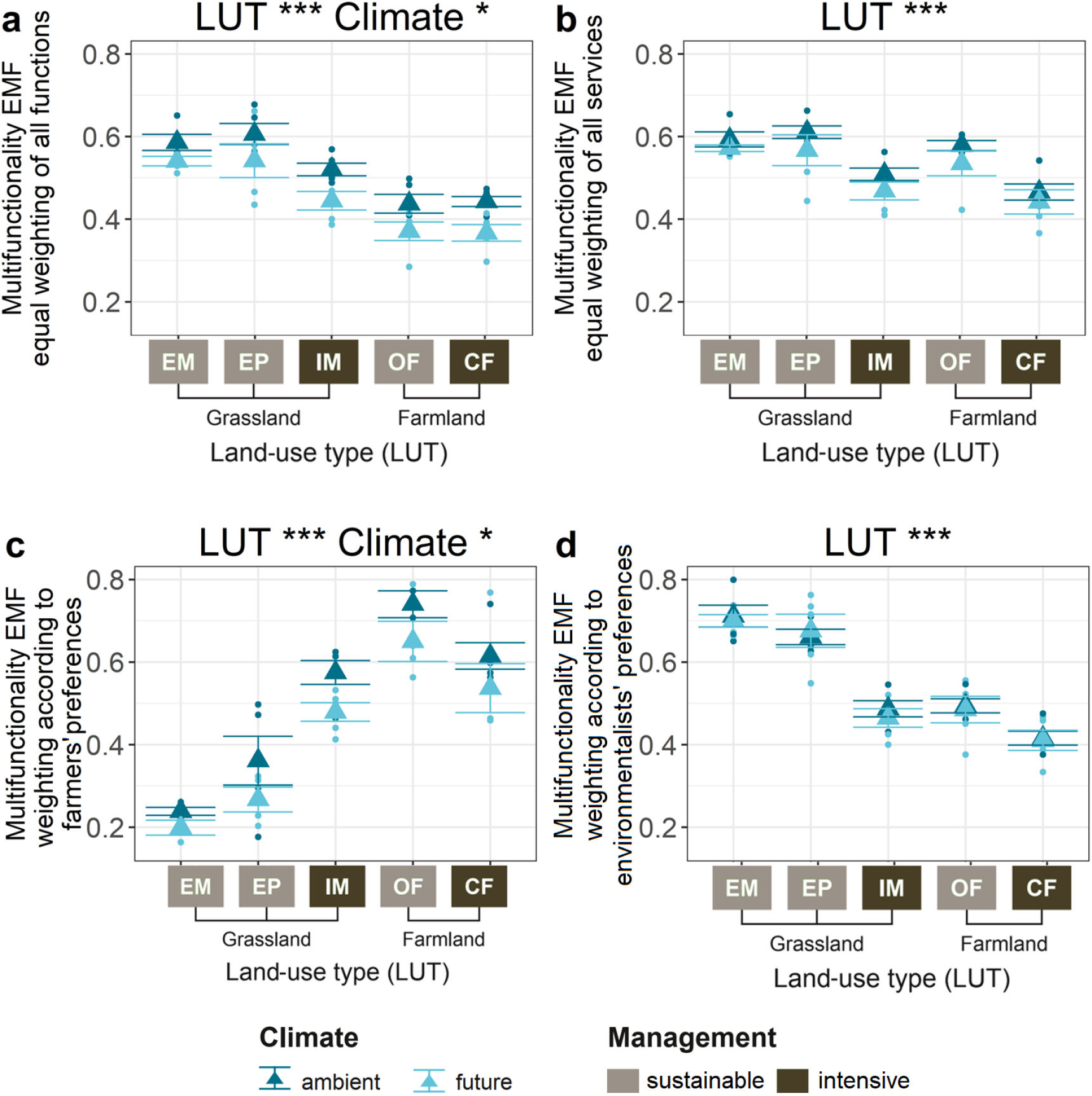
Ecological ecosystem multifunctionality as affected by five different land-use types (EM: extensive meadow, EP: extensive pasture, IM: intensive meadow, OF: organic farming, CF: conventional farming) for both the ambient and the future climate and for four different weighting scenarios (a: equal ecosystem function weighting, b: equal ecosystem service weighting, c: weighting according to farmers’ preferences, d: weighting according to environmentalists’ preferences). Dots indicate the multifunctionality level within the plots of the experiment, triangles indicate group means. Asterisks indicate a significant effect of the respective factor or interaction (*** p<0.001; ** p<0.01; * p<0.05).

### 3.2. Effects of climate and land-use type on economic multifunctionality measures

Considering the economic measure of EMF value, future climate does not show a significant effect. However, the land-use type has a strong impact on economic EMF (p<0.001***; Figure 4). The total economic value of ecosystem service provision is, on average, higher for grassland compared to farmland. For both grassland (p<0.01**) and farmland (p<0.001***), economic EMF, which includes economic benefits to society at large, is around ∼1.7 times higher for sustainable compared to intensive management. For sustainably-managed grassland, economic EMF is around twice as high as the monetary ecosystem service provision for farmers (consisting of the ES food production, biodiversity conservation and soil health), while both values are similar for intensively-managed grassland and sustainably-managed farmland. For intensively-managed farmland, monetary ecosystem service provision for farmers is 1.42 times higher than the total economic EMF value (see Figure 3). Economic EMF composition differs between grassland and farmland, as well as between sustainable and intensive management. For grassland, a significant proportion (39 to 62%) of the economic EMF value is contributed by the service climate regulation. Under intensive management, the economic value of food production is slightly increased for grassland, and slightly decreased for farmland. For the intensively-managed land-use types, nitrogen surplus has a strongly negative impact on economic EMF, an effect that is negligible for the sustainably-managed types. The monetary value of soil health is dependent on the level of biodiversity in the respective plot, and *vice versa*, due to the mutual effect of soil health and biodiversity on yield stability.

**Figure 3.**
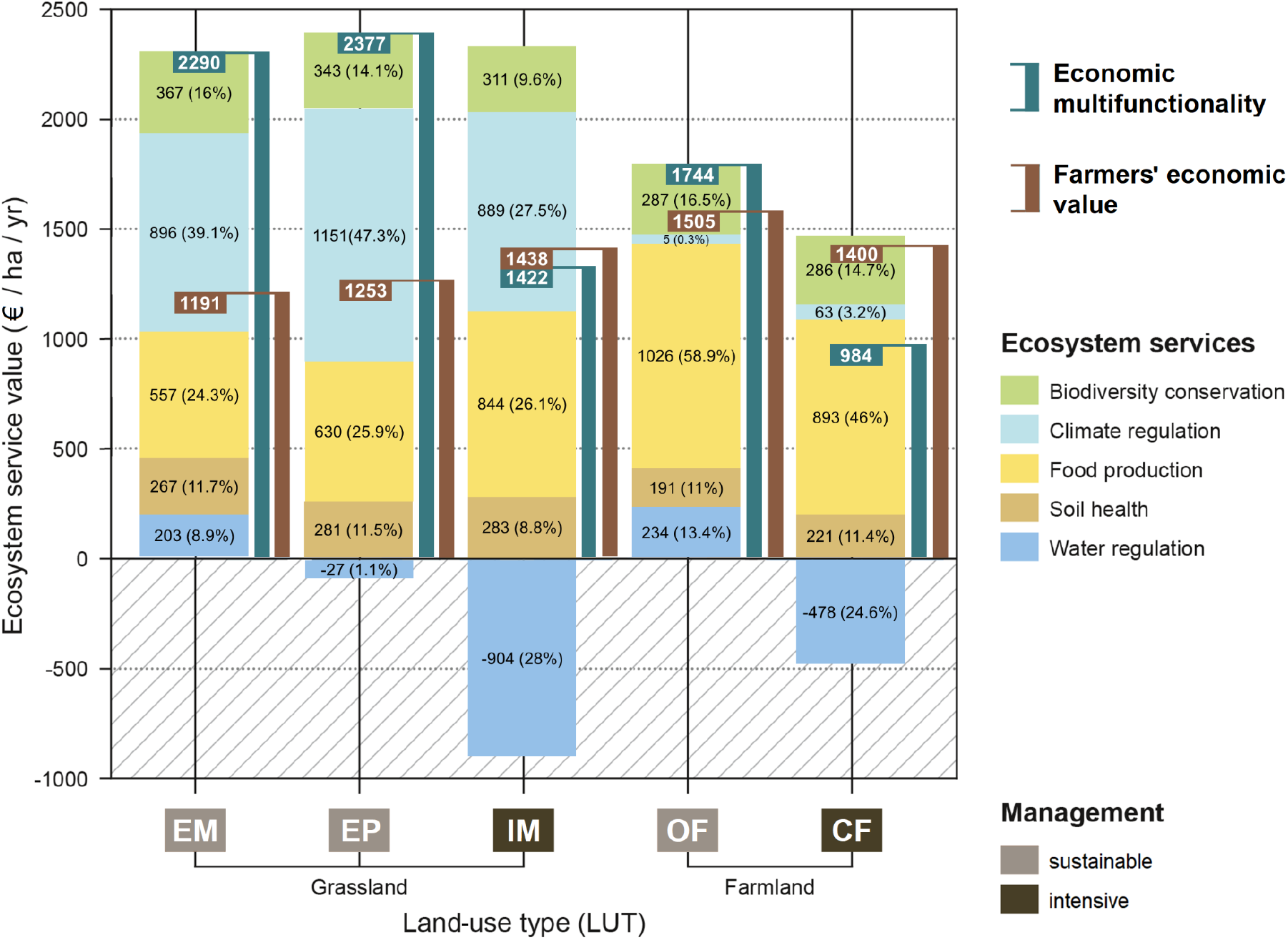
Yearly ecosystem service provision in monetary units for different land-use types (EM: extensive meadow, EP: extensive pasture, IM: intensive meadow, OF: organic farming, CF: conventional farming). Vertical bars indicate the economic ecosystem multifunctionality value (eEMF, blue, including benefits for society at large) and farmers’ economic value (brown, composed of food production and yield stabilising services biodiversity and soil health).

**Figure 4.**
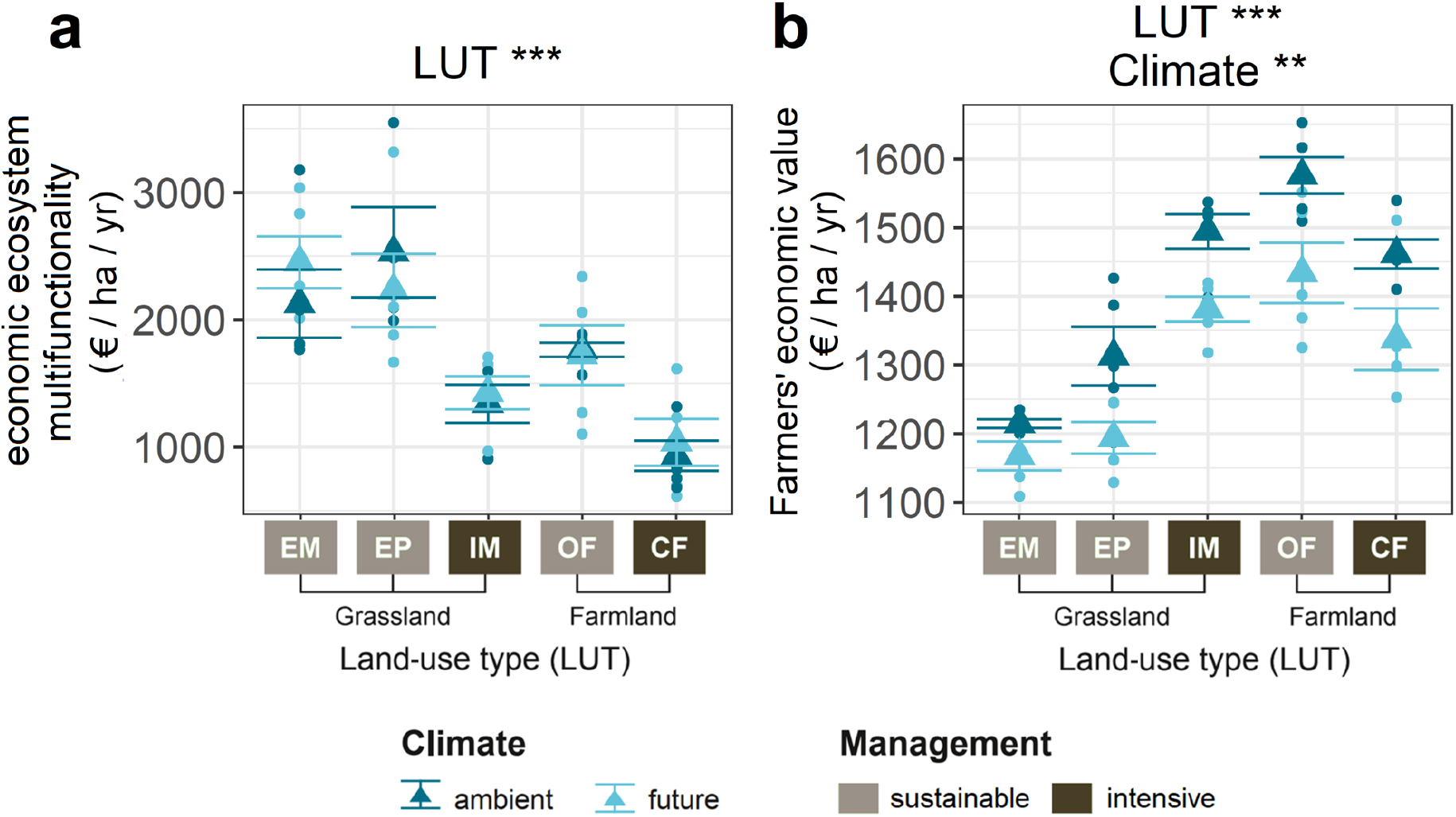
Economic ecosystem multifunctionality value (eEMF, a) and farmers’ economic value (FEV, b) as affected by five different land-use types (EM: extensive meadow, EP: extensive pasture, IM: intensive meadow, OF: organic farming, CF: conventional farming) for both ambient and future climate. Dots indicate the level of the economic multifunctionality and farmers’ economic value within the plots of the experiment, triangles indicate group means. Asterisks indicate a significant effect of the respective factor or interaction (*** p<0.001; ** p<0.01; * p<0.05).

With a mean of 2024 € / ha (grassland) and 1364 € / ha (farmland), the yearly economic value of EMF is substantially higher than the land rent for land of similar quality in the region where the experiment we obtained our data from is located (348 € / ha / yr for grassland, 574 € / ha / yr for farmland in the Saalekreis district 2019) (Ministry for the Environment, Agriculture and Energy of the State of Saxony-Anhalt, 2020). While economic EMF is only affected by land-use type, farmers’ economic value is also significantly decreased under future climate (see Table 3, Figure 4).

**Table 3.** General linear regression model table of F and p values (in brackets: numerator and denominator d.f.) of the effect of the two factors land-use type (LUT), climate type, and their interaction on economic ecosystem multifunctionality value (eEMF) and on farmers’ economic value (FEV). Bold values indicate a significant effect of the respective factor or interaction (*** p<0.001; ** p<0.01; * p<0.05).

|  | LUT |  |  | Climate |  |  | LUT:Climate |  |
| --- | --- | --- | --- | --- | --- | --- | --- | --- |
| Value | F(4,32) | p value |  | F(1,8) | p value |  | F(4,32) | p value |
| eEMF | **19.41** | **0.000** | *** | 0.06 | 0.81 |  | 0.70 | 0.60 |
| FEV | **49.56** | **0.000** | *** | **17.65** | **0.003** | ** | 0.96 | 0.441 |

**Table 4.** General linear regression model table of F and p values (in brackets: numerator and denominator d.f.) of the effect of the two factors land-use type (LUT), and climate type (Climate), and their interaction (LUT:Climate) on each individual ecosystem function. Bold values indicate a significant effect of the respective factor or interaction (*** p<0.001; ** p<0.01; * p<0.05).

|  | LUT |  |  | Climate |  |  | LUT:Climate |  |  |
| --- | --- | --- | --- | --- | --- | --- | --- | --- | --- |
| Ecosystem function | F(4,32) | p value |  | F(1,8) | p value |  | F(4,32) | p value |  |
| Yield | **97.22** | **0.000** | *** | **8.12** | **0.021** | * | 0.57 | 0.684 |  |
| Total organic soil carbon | **15.60** | **0.000** | *** | 3.96 | 0.082 |  | 0.18 | 0.950 |  |
| Nitrogen surplus | **213.66** | **0.000** | *** | **5.72** | **0.044** | * | 0.32 | 0.860 |  |
| Microbial Biomass | **37.82** | **0.000** | *** | 0.11 | 0.749 |  | **3.21** | **0.025** | * |
| Cellulase activity | 1.68 | 0.179 |  | **5.34** | **0.050** | * | 0.24 | 0.913 |  |
| N-acetylgluco-saminidase activity | **22.13** | **0.000** | *** | 1.51 | 0.254 |  | 0.64 | 0.639 |  |
| Acid-phosphatase activity | **7.02** | **0.000** | *** | 1.38 | 0.274 |  | 1.18 | 0.337 |  |
| Belowground decomposition rate | **8.02** | **0.000** | *** | 1.63 | 0.237 |  | 0.83 | 0.514 |  |
| Aboveground decomposition rate (microbes) | **22.67** | **0.000** | *** | 0.08 | 0.790 |  | 0.86 | 0.497 |  |
| Aboveground decomposition rate (microbes + fauna) | 0.67 | 0.617 |  | 1.92 | 0.173 |  | 0.64 | 0.639 |  |
| Biodiversity (mesofauna) | 2.64 | 0.052 |  | 3.90 | 0.084 |  | 0.21 | 0.933 |  |
| Biodiversity (macrofauna) | **23.80** | **0.000** | *** | 1.07 | 0.331 |  | 1.24 | 0.312 |  |
| Biodiversity (nematodes) | **16.28** | **0.000** | *** | 3.90 | 0.055 |  | 0.88 | 0.485 |  |

### 3.3. The role of biodiversity for ecosystem multifunctionality

The multiple threshold approach was used to analyse the biodiversity-multifunctionality (EMF) relationship (Byrnes et al., 2014). Therefore, EMF was composed of only 10 ecosystem functions, whereas biodiversity-related functions were not included, but used to calculate an overall biodiversity metric (calculated as multidiversity to integrate biodiversity data of different taxonomic groups) that served as explanatory variable. Here, a positive correlation between biodiversity and EMF was observed (R^2^ = 0.246, p<0.001***). A display of the slope of the regression line of the biodiversity-EMF relationship as function of multiple thresholds for both ambient and future climate reveals insights into the nature of the biodiversity-EMF relationship by comparison of the characteristic graph points T_min_ / T_max_ (smallest / highest threshold at which biodiversity starts to affect EMF) and T_mde_ (threshold where biodiversity has the strongest effect on EMF) (Byrnes et al., 2014). No significant difference was found regarding the strength of the biodiversity-EMF relationship under ambient or future climate (T_min_ at threshold t = 29% for both ambient and future climate, T_max_ at t = 71% for both ambient and future climate, T_mde_ at t_amb_ = 51% / t_fut_ = 49%) (see Figure 5).

**Figure 5.**
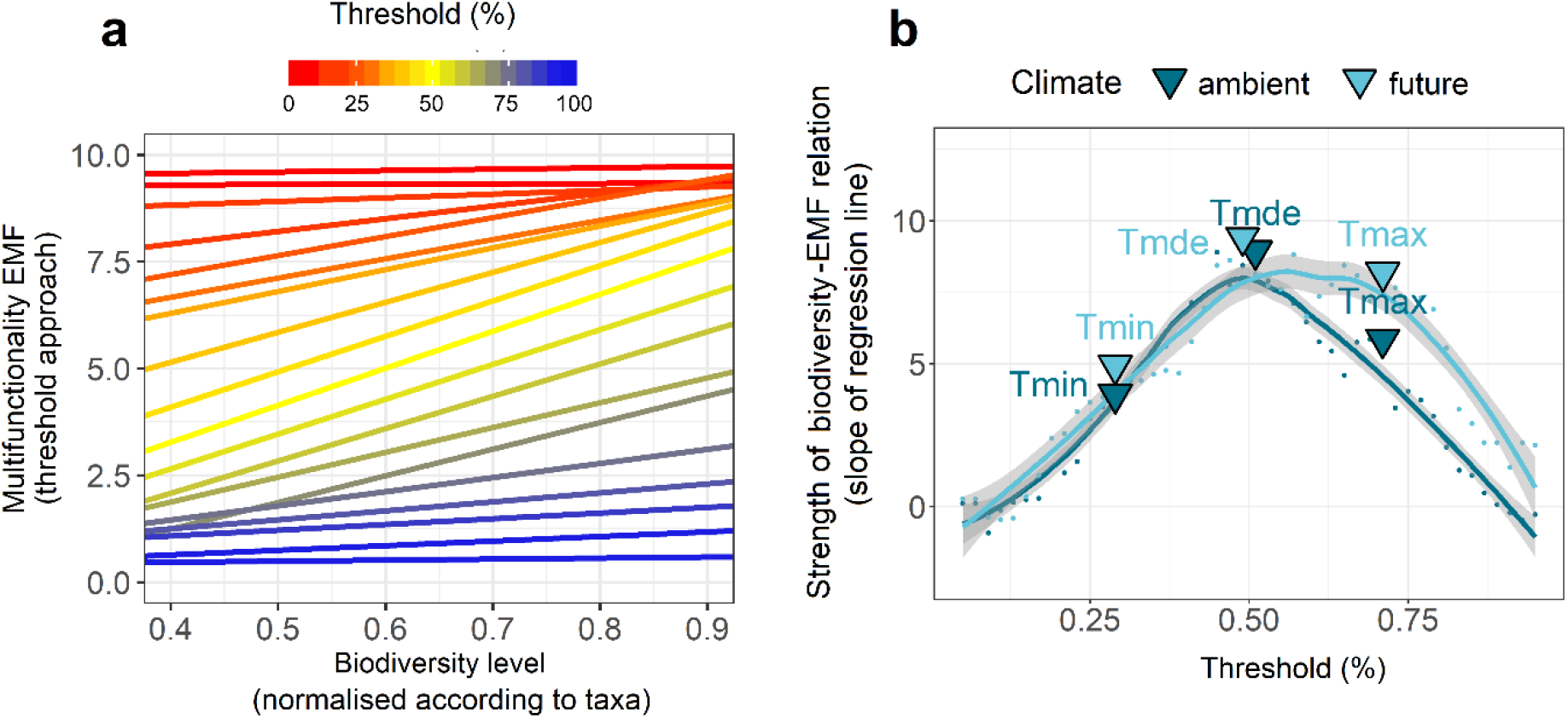
Analysis of the biodiversity-multifunctionality relationship using the multiple threshold approach (biodiversity was calculated as multidiversity). (a) Regression lines for the biodiversity-multifunctionality relationship for different thresholds between 0 and 100% including data from both ambient and future climate. (b) Strength of the biodiversity-multifunctionality relationship (slope of the regression line) as a function of different thresholds for both ambient and future climate. Characteristic graph points that were used to analyse differences in the biodiversity-multifunctionality relationship under the two climate types are marked.

**Figure 6.**
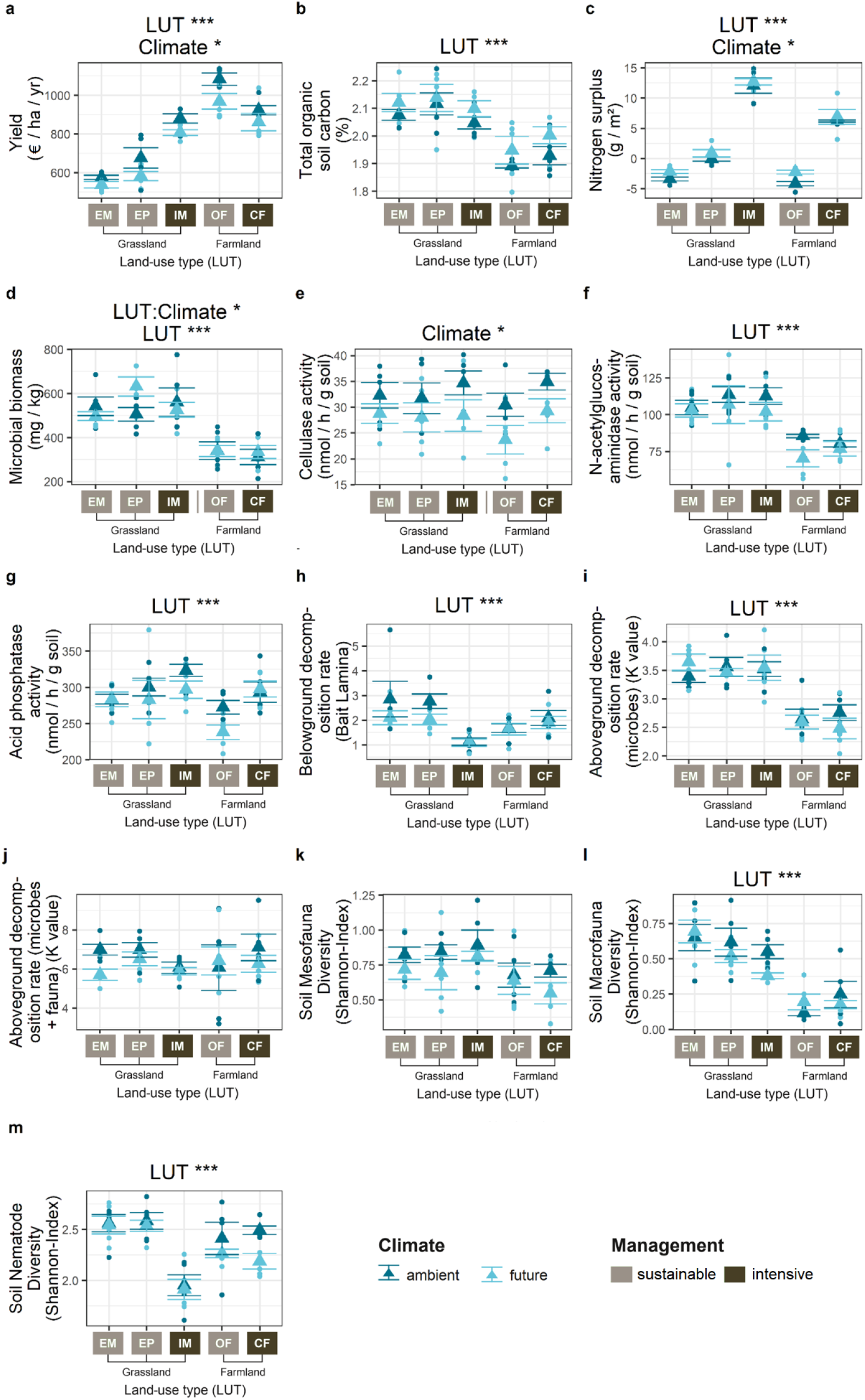
Individual ecosystem function levels as affected by five different land-use types (EM: extensive meadow, EP: extensive pasture, IM: intensive meadow, OF: organic farming, CF: conventional farming) for both ambient and the future climate. Dots indicate the ecosystem function levels within the plots of the experiment, triangles indicate group means. Asterisks indicate a significant effect of the respective factor or interaction (*** p<0.001; ** p<0.01; * p<0.05).

## 4. Discussion

The present study shows that the effects of future climate and land-use type on ecosystem multifunctionality (EMF) are strongly dependent on the type of multifunctionality measure. We contrasted ecological EMF measures, based on averaging approaches with different weighting scenarios of ecosystem functions, to an economic EMF measure based on the value of multiple ecosystem services provided to farmers and to society at large. Future climate showed significantly adverse effects on EMF for an equal weighting of all functions and for the weighting according to farmers’ preferences. However, the expected net negative effect of future climate on EMF under an equal weighting of all ecosystem services (Scholes 2016) was not supported by the data in hand. The rather weak effect of the future climate type might be partially explained by the relatively mild temperature treatment in the GCEF with an increase of about +0.55°C which is in the lowest range of predictions at the time of the establishment of the experiment. Contrary to projections a decade ago, this future climate treatment may be a conservative estimate (Korell et al., 2020) of alterations towards the end of this century, with more recent predictions ranging between +1.1°C to +5.5°C, (reflecting all possible scenarios) and a mean of +3°C (Regional Climate Atlas Germany, 2021). The GCEF is among the few infrastructures which manipulate precipitation according to a realistic mean future scenario (Korell et al., 2020). However, it can be assumed that the applied projected mean change of climatic conditions will have less severe effects on agroecosystems than the increased frequency of climate extreme years which is an concomitant phenomenon of future climate. As a consequence, we expect that future climate change may have even stronger effects on EMF than presented in this study.

The present study further suggests that future climate effects are consistent across land-use types. This finding does not support our hypothesis of significant differences in the resilience of certain land-use types towards the effect of future climate (Isbell et al., 2015; Tilman & Downing, 1994). EMF of all land-use types was equally negatively affected (or not significantly affected) by future climate which contradicts other research showing a substantial spatial heterogeneity in sensitivity of ecosystem service provision to future climate (Hua et al., 2021; Scholes, 2016; IPCC, 2014) but goes align with findings showing grasslands being resilient towards future climate (severe droughts) regardless of the management intensity (Buchmann et al., 2019). Contrary to future climate, land-use type and management regime could be shown to cause pervasive effects on multiple ecosystem functions and on EMF in this study.

From a farmer’s perspective, EMF is highest for the intensively-managed grassland type, and for both farmland types, as these land uses maximise the ecosystem service ‘yield’, which has the highest weighting assigned here, whereas the dis-service nitrogen surplus (which is high for the intensively managed land-use types) is hardly considered from a farmer’s perspective. The sustainably-managed farmland type reaches the highest yields, as organically produced crops obtain high prices (subsidies not included), which outweigh the slightly lower mass yields (7%) compared to conventional crops, an effect also shown before (Wittwer et al., 2021). This calculation does not consider that a shift to organic farming might change costs (reduced expenses for agrochemicals) and effort (e.g., mechanical weeding) for farmers. Moreover, yields will further depend on the sown crop species. Further, due to the monetisation based on current market values, the yield data does not represent the intrinsic value of the ecosystem. Still, there was no other meaningful way to make yields of a large number of different crops comparable. Keeping this in mind will help to draw reasonable conclusions from the results of this study.

Under a farmer’s perspective, EMF (which can be seen as the multidimensional benefit farmers receive from ecosystems) is strongly decreased under future climate. This is explained by the role of water as a limiting factor for plant growth during the growing season aligning with predictions of a net-negative effect of future climate on agricultural production for Europe (IPCC, 2019; Olesen & Bindi 2002). By that, we expect farmers to be severely affected by climate change if adequate adaptation measures are not taken.

From a perspective of environmentalists, the intensively managed land-use types are less valued. This is partly due to their high level of nitrogen surplus due to the application of mineral nitrogen fertiliser. During the growing season, nearly all available mineral nitrogen is assimilated by the plants if the conditions are suitable for plant growth. Contrary, in periods of drought, plant growth can stagnate despite the addition of mineral nitrogen fertiliser. The lack of nitrogen uptake by plants during such periods increases the chances that unused nitrogen surplus leaches to water bodies and groundwater when sufficient water is available outside the growing season (Klaus et al., 2018). This reduced plant uptake might explain the increased nitrogen surplus under future climate. Due to the high soil water holding capacity, a deep root penetration and low precipitation within the area of the experiment (Schädler et al., 2019), it remains unclear if nitrogen leaching occurs or whether the specific site conditions allow for complete denitrification.

From the perspective of environmentalists, not only the intensively-managed land-use types but also the sustainably-managed farmland performs poorly due to its low levels of several ecosystem functions that contribute to soil health (microbial biomass, aboveground decomposition rate; Guerra et al., 2021; Wall et al., 2015). Here, the sustainably-managed grassland types are clearly preferred, as they show no nitrogen surplus and high levels in soil health related ecosystem functions. This finding is in line with previous work showing that species-rich grasslands reduce nitrogen leaching from soils (Leimer et al., 2016), support higher soil microbial biomass, activity, and diversity (Eisenhauer et al., 2010; Lange et al., 2015), and elevated decomposition rates (Vogel et al., 2013).

As an alternative to ecological EMF measures based on averaging approaches, we proposed an economic measure based on the economic values of various ecosystem services. This economic EMF measure includes both values for the farmer (e.g. yield and insurance values) and for society at large (e.g. carbon sequestration). In general, grasslands showed a higher economic EMF value than farmlands, and for both grassland and farmland, the sustainably-managed types showed values ∼1.7 times higher than the intensively-managed types. Despite the expectation of a decrease in the ecosystem service provision under future climate (IPCC, 2019; Scholes, 2016), no significant decrease through future climate was shown here (3% decrease based on comparison of mean values). This suggests that sustainable land use is a promising approach to maintain and promote the economic EMF value of managed land also under a changing climate.

The value composition of the five land-use types differed strongly though. For grassland, the value for food production is slightly increased under intensive management due to the application of fertiliser. For farmland, sustainable management reaches higher monetary yields due to the higher sales prices of legumes and organically-produced crops, as shown before (Wittwer et al., 2021). The benefits in productivity in intensively-managed grassland are outweighed by the disadvantages of increased nitrogen surplus that significantly decreases the economic EMF value of all intensively-managed land-use types. This finding is of high relevance, given that surplus nitrogen and nitrogen contaminating groundwater is a critical issue in many intensively-managed regions of the world (Lin et al., 2001; Singh & Sekhon, 1979) and represents a threat to human health (Ahmed et al., 2017; Majumdar & Gupta, 2000; Wolfe & Patz 2002).

For grassland, a significant proportion (39 to 62%) of the economic EMF value is contributed by the ecosystem service ‘climate regulation’, due to the carbon storing effect in the soil (∼10% increase in total organic carbon over the first three years after the implementation of the treatments), which indicates the economic potential for land managers (*carbon farming*) under an externality rewarding regime. As the site of the experiment was used as farmland before the implementation of the treatments, the carbon storage effect is negligible for farmland types, as they might have already reached their soil carbon equilibrium and their yearly carbon flows are about zero. Due to the incremental increase of the soil carbon content over years (Lange et al., 2019) and its high interconnection with other ecosystem functions (Smith et al., 2021; Lange et al., 2019), land-use change requires time to affect EMF. By that, the detrimental effects of intensive land-use might increase in the future due to a loss of soil communities and soil related ecosystem functions.

The monetary values of soil health and biodiversity are based on their stabilising effect on yield, which aligns with previous research (Dasgupta, 2021; Gao et al., 2015; Watts & Torbert, 2011; Zhang et al., 2009). This insurance value of biodiversity and soil health is dependent on the risk aversion and might be smaller for less risk averse stakeholder(s) (see Figure 9, Supplementary material). Moreover, different findings of this study suggest that biodiversity is an important driver of ecosystem multifunctionality, such as indicated by a significant positive correlation between soil biodiversity and EMF (equal ecosystem function weighting) (R^2^ = 0.247, p<0.001***). Although the experimental design does not allow proofing a causal relation, this observation goes in line with former research showing soil biodiversity to be a main driver of EMF (Delgado-Baquerizo et al., 2020; Schuldt et al., 2018; Delgado-Baquerizo et al., 2016; Soliveres et al., 2016; Wagg et al., 2014). For the grassland types, for instance, the low multifunctionality level of the intensively-managed type goes hand in hand with a low (nematode-)biodiversity. This further aligns with the findings showing that biodiversity loss is a main driver of ecosystem multifunctionality reduction due to land-use intensification (Allan et al., 2015). The positive relationship between biodiversity on EMF was equally strong under ambient and under future climate in the present experiment. While this aligns with the findings describing an increase of (soil) multifunctionality with higher (plant) biodiversity in both ambient and future environments (Eisenhauer et al., 2018), it contrasts other findings describing a stronger effect of biodiversity on ecosystem functioning in more stressful environments (Hong et al., 2022).

Taken together, the present work clearly highlights the risk of a significant decline in EMF due to land-use intensification, climate change, and biodiversity loss which has potentially adverse consequences for humans. Therefore, sustainable land management that shows the highest level of EMF should be promoted as well as measures and incentives implemented to increase biodiversity within agricultural areas fostering EMF. Notably, this study also introduces an important conflict. Agricultural land is typically managed by farmers. However, those are the only stakeholder group considered in this study, whose multidimensional benefit obtained from EMF was higher under intensive management (for grassland types) – at least in the short term –, which is underlined by the value gap between farmers’ monetary benefit received from agroecosystems and their respective economic EMF value for society. As a consequence, we suggest to provide incentives to make farmers choose the land management that is preferred from a macrosocial perspective and to adapt compensation schemes that currently put too little emphasis on sustainable management practice and environmental measures, working towards an economy incorporating external costs.

## 5. Materials and methods

### 5.1. Experimental setup

Data was obtained from the Global Change Experimental Facility at the field research station of the Helmholtz Centre for Environmental Research in Bad Lauchstädt, Saxony-Anhalt, Germany (51°22’60 N, 11°50’60 E, 118 msl) (Schädler et al., 2019). This area is characterised by a sub-continental climate and predominantly west winds with mean annual precipitation of 489 mm (1896-2013), respectively 525 mm (1993-2013), and mean temperature of 8.9°C (1896-2013), respectively 9.7°C (1993-2013). The soil of the study site is a Haplic Chernozem representing one of the most fertile soils to be found in Germany (Schädler et al., 2019). The GCEF was designed for a simultaneous manipulation of land-use type (5 types: 2 farmland and 3 grassland types) and climate (ambient and future), using a fully randomised split plot experimental design that allows full-factorial combination of the climate and land-use types with 50 plots of approximately 400 m^2^ each (Schädler et al., 2019). The experiment was established in 2013 on a field formerly used as farmland and has been ongoing since then. Manipulation of climate and the establishment of land-use types started in 2014. Land-use types represent five agricultural management forms typically practiced in Germany: Conventional farming (CF) including a crop rotation with winter rape, winter wheat and winter barley and application of mineral fertilisers and pesticides; organic farming (OF) including legumes in the crop cycle every three years (alternating alfalfa and white clover) to replace mineral N fertilizer as well as only physical weed control without the application of herbicides; intensive meadow (IM) that includes the sowing of a seed mixture (four different *Poaceae* species) and the application of mineral fertiliser at the beginning of the growing season as well as after the first, second and third cut; extensive meadow (EM) that includes the sowing of a seed mixture with 56 species (14 grass species, 10 legumes and 32 herbs) representing all forms of vegetable life of grassland typically in Germany; and extensive pasture (EP) that includes the same seed mixture as EM and a grazing with sheep (∼20 sheep grazing 24 hours per plot) which takes place three times a year (early spring, mid- to late spring, and mid of summer). The land-use types can be differentiated into grassland (IM, EM, EP) and farmland (OF, CF), and into sustainable (OF, EM, EP) and intensive (CF, IM) management. Sustainable management refrains completely from the application of agrochemicals (mineral nitrogen fertiliser and pesticides). Climate manipulation is reached with large steel constructions that cover each plot, equipped with mobile shelters, side panels and an irrigation system, whereas night temperature is passively increased via automatic closing of the shelters and panels from sunset until sunrise resulting in an 0.55°C increase of daily mean temperature and a stronger increase by up to 1.14°C in average in minimum temperature. Summer precipitation is reduced by ∼20% via control of the roofs by a rain sensor. Precipitation in spring and autumn is increased by ∼10% with an irrigation system. With this treatment, a consensus scenario across different models of climate change in Central Germany for the years between 2070 and 2100 is simulated. The control plots that are managed under ambient conditions are equipped with the same steel construction to exclude possible microclimate effects on the experimental setup. Before the start of the experiment, oat was sown on all subplots in 2013 to homogenise soil conditions (Schädler et al., 2019).

### 5.2. Samplings and measurements

During the years 2014 to 2020, plots were harvested with a combine harvester. Yield biomass (dt / ha, for farmland differentiated into grain and straw yields) was measured after air drying which left the biomass with residual moisture of 14% (barley / wheat grains) and 9% (rape grains). Depending on the annual environmental conditions, for grassland, harvesting occurred up to four times per year. For the total productivity over the year, yields of all harvests are summed up for each plot. For extensive pasture, machine harvest was not practical, as plots were grazed with sheep. Instead, harvesting was done manually right above the soil (for each plot, four subsamples were taken and averaged). Here, total yield also considers the grazing uptakes of the sheep, measured as the difference between the biomass in four subsamples in sheep-excluding cages and four subsamples in the sheep area. (Bio-)mass yield was converted to monetary yield based on producer prices (Table 5, Supplementary material).

**Table 5.**
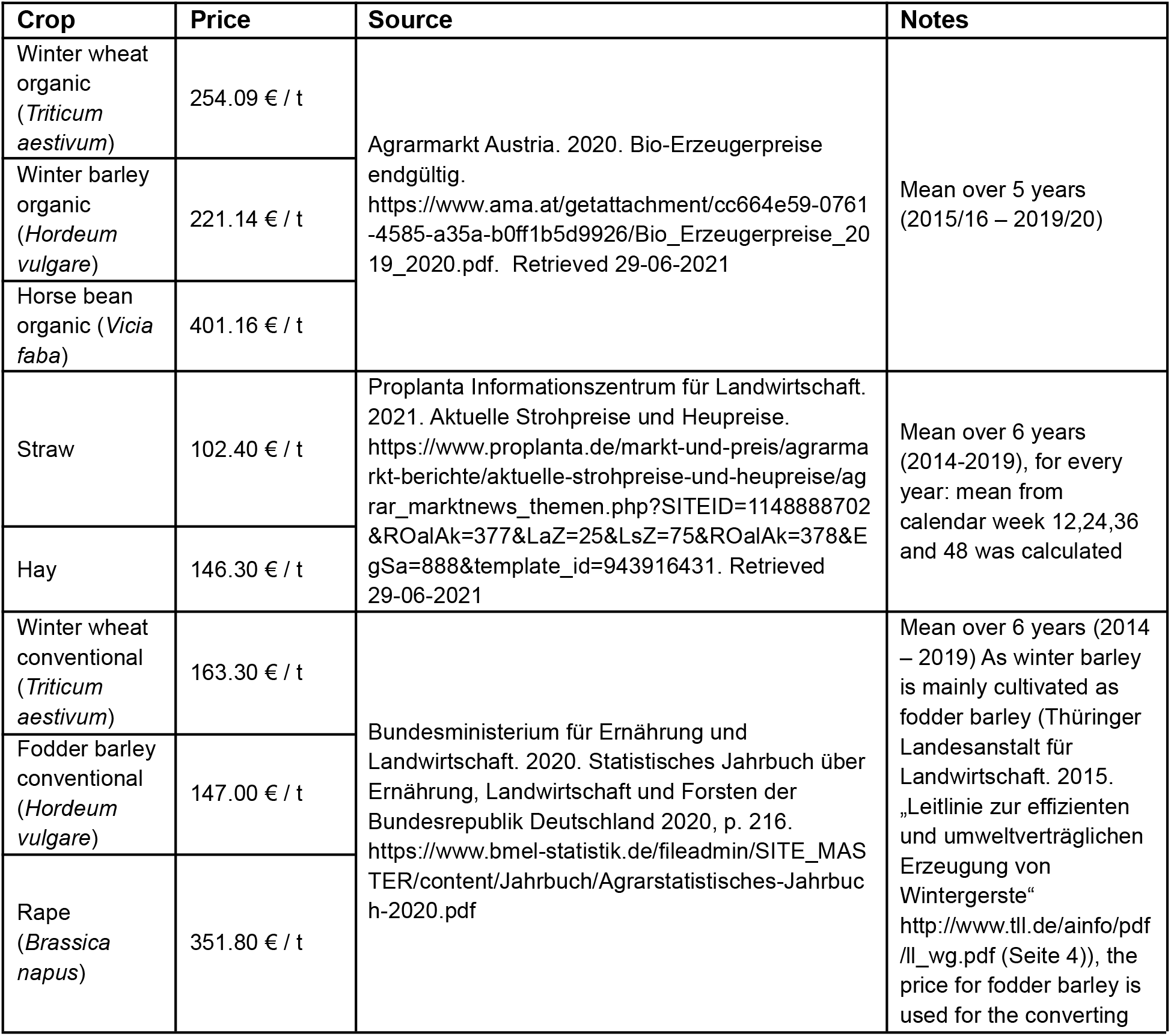
Producer prices used for the conversion of the yield unit from dry biomass into monetary value with information regarding source and conversion process.

Soil samples were taken annually in the beginning of September (mean of 5 measurements per plot) to measure total organic soil carbon (% of dry soil, measurement according to Breitkreuz et al. (2021)), microbial biomass that can be used as a proxy for the soil water holding capacity which is (next to nutrient supply) the most important factor defining plant growth as microbes require an aquatic medium to survive and proliferate (μg microbial biomass carbon / g dry soil, measurement according to Francioli et al. (2016)), enzymatic activity (nmol / h / g dry soil, measured at pH 5 according to Francioli et al. (2016), three enzymes that are ubiquitous in most organisms were measured as indicator for the rate at which microbes can decompose and process organic matter to provide nutrients that are accessible to plants (phosphorous, nitrogen)). Belowground decomposition rate that is closely linked to the biogeochemical nitrogen cycle and includes effects of large decomposers and fungi was measured 3-weekly from 2015 to 2016 using bait lamina measurements (average of 6 bait lamina stripes within one plot that remained in the soil for 3 weeks). Aboveground decomposition rate (microbes / microbes + fauna) was measured using litterbags (0.02mm / 5mm mesh size) left on soil for two (summer and spring) or 4 months (winter) with a total of 7 measurement periods, starting in April 2015 (see Yin et al. (2019a) for methodological details). Soil mineral nitrogen (NH_4_ ^+^ and NO_3_ ^-^) content (mg / kg dry soil) was measured in 3-weekly resolution from 2015 to 2017 according to Breitkreuz et al. (2021). Mineral nitrogen deprivation (kg N / ha) was measured over the years 2016 to 2019 through an elemental analysis of the harvest (for organic and conventional farming only for the machine harvest (cut 10 cm above soil), for intensive and extensive meadow also for the manual harvest (cut 3 cm above the soil surface), for extensive pasture only for the manual harvest). Plant material (dried at 70°C for 48 hours) was shredded and homogenised, a subsample was milled to a fine powder, and appr. 10 mg of the finely milled plant material were weighted with an analytical microbalance (Cubis MSA 3.6P, Sartorius AG, Göttingen, Germany) into tin capsules and measured with an elemental analyser (Vario EL cube, Elementar Analysensysteme GmbH, Langenselbold, Germany). Nitrogen stocks were calculated based on data on yield (dry biomass). Soil biodiversity (Shannon-Index) was measured from years 2015 to 2016 (two measurements per year (spring and autumn), for methodological details see Yin et al. (2019b) (meso- and macrofauna) and Siebert et al. 2020 (nematodes)).

### 5.3. Calculation of nitrogen surplus

Nitrogen surplus was calculated on plot level adapted from Richner et al. (2014) as the soil mineral nitrogen content at time x plus the annual input of mineral nitrogen (Schädler et al., 2019) minus the annual nitrogen deprivation through the harvest plus the soil mineral nitrogen content at time x + 1 year. For x, the time between one year’s harvest and the sowing / first fertilisation for the next year’s harvest was chosen (end of July to beginning of August for farmland types; end of January to beginning of February for grassland types). The mean of two measures was used due to the variability of the nitrogen state due to weather conditions. Different datasets were unified (mineral nitrogen content in g / m^2^ and in mg / kg soil) based on the assumption that the applied fertiliser spreads and accumulates in the upper 20 cm of the soil and that soil weight is 1350 kg/m^3^ (Engineering ToolBox, 2008). Nitrogen outgassing and deposition were not considered here as they are opposing processes that can equalise each other (Richner et al., 2014).

### 5.4. Calculation of multidiversity

To integrate different datasets on biodiversity (or, biodiversity levels of different organism groups) into a proxy for the overall biodiversity level, the “multidiversity” was calculated as the average proportional biodiversity across the taxonomic groups, (or respectively, size categories that usually contain certain taxonomic groups), normalised with the maximum observed level of biodiversity of the respective dataset (Allan et al., 2014).

### 5.5. Calculation of soil health

Soil health was calculated as the average of the normalised level of the three ecosystem functions microbial biomass, enzymatic activity (calculated as the average of the normalised performance of the three ecosystem functions cellulase activity, N-acetylglucosaminidase activity and acid-phosphatase activity) and decomposition rate (calculated as the average of the normalised level of the three ecosystem functions belowground decomposition rate, aboveground decomposition rate (microbes) and aboveground decomposition rate (microbes + fauna)).

### 5.6. Calculation of ecosystem multifunctionality

To calculate ecosystem multifunctionality, two ecological approaches are used to address different aspects of the research objectives. Within the **averaging approach**, ecosystem multifunctionality EMF is calculated as weighted average of the levels of the different ecosystem functions *EF*_*i*_ with *α* as a weighting factor, where *EF*_*i*_ is calculated as a fraction of an actual value *X*_*i*_ to two reference values (minimum (*X*_*i,min*_) and maximum (*X*_*i,max*_) observed value of the respective ecosystem function *i*) (Byrnes et al., 2014).

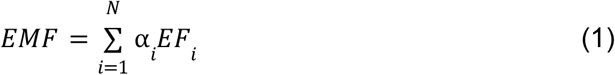

To calculate EMF according to different stakeholders’ perspectives and objective scenarios, ecosystem functions and services were weighted according to different objectives and stakeholder priorities (stakeholder priorities adapted from Manning et al., 2018, see Figure 7, Supplementary material). Equal service weighting and weighting according to farmers’ preferences give less weight to ecosystem functions related to soil health and biodiversity. Further, farmers assign less weight to regulating services but give high weight to food production, which is not considered in the weighting according to environmentalists’ preferences.

**Figure 7.**
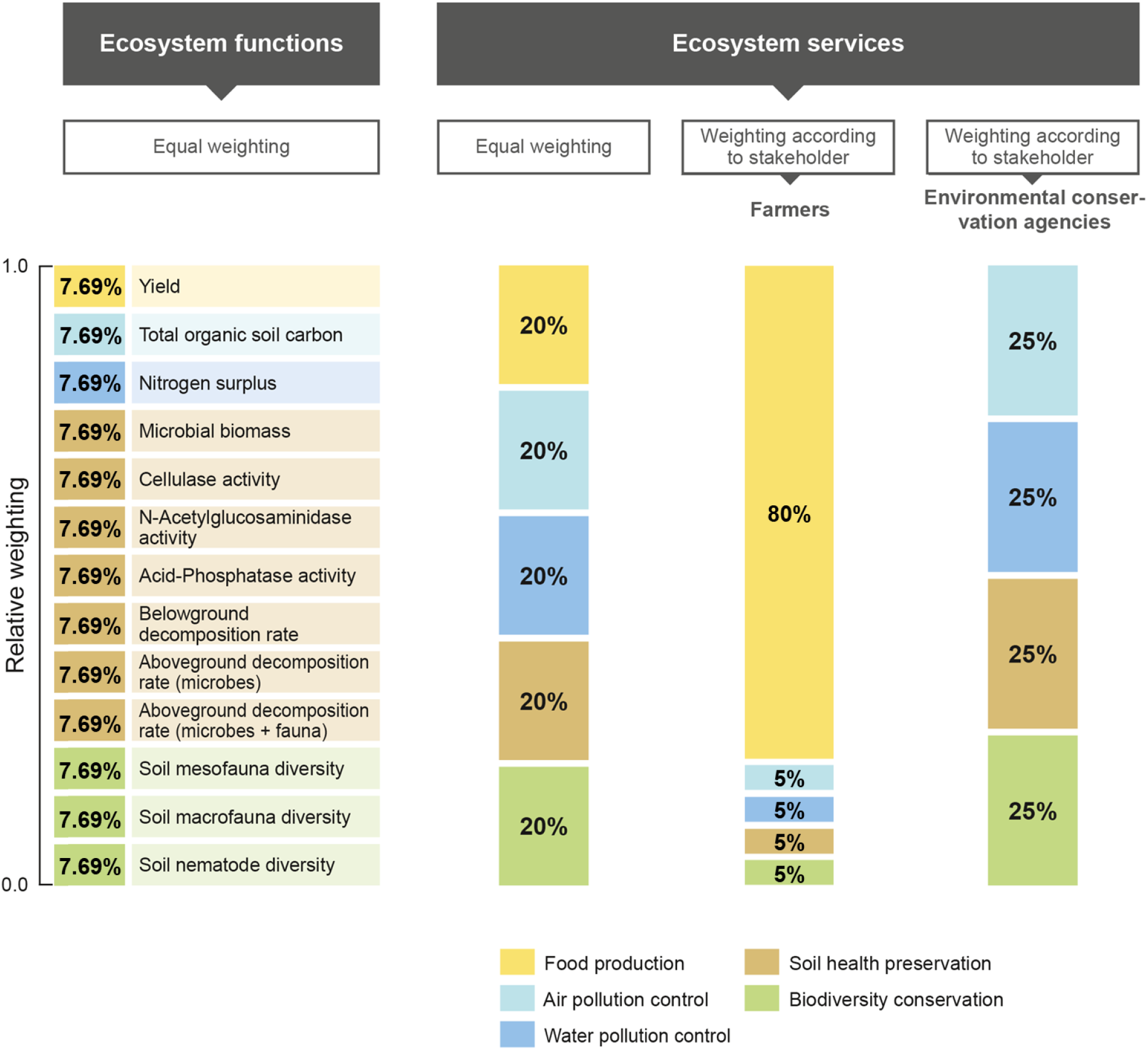
Weighting of the 13 ecosystem functions (or 5 ecosystem services, respectively) within four different multifunctionality scenarios.

To investigate biodiversity-multifunctionality relationships, the multiple threshold approach was used (Byrnes et al., 2014). Here, the number of functions *EF*_*i*_ that perform above a certain threshold *t* (fraction of the maximum value of the respective function) represent multifunctionality *EMF* within the respective plot. With this approach, we account for the nature of ecosystem multifunctionality, whereas a loss of function or service delivery can be accepted until a certain degree (threshold), under which the function of the respective function or service is lost.

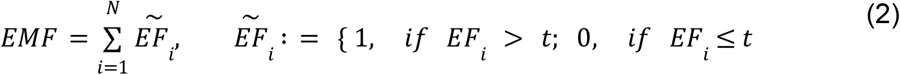

The application of multiple thresholds within the range of 0% and 100% reveals key information about the relationship of biodiversity and ecosystem multifunctionality, by plotting the slope of the biodiversity-multifunctionality regression line against the range of thresholds (Byrnes et al., 2014).

### 5.7. Quantification of economic ecosystem multifunctionality value

Yield raw data was obtained as data on dry biomass and monetised based on current market prices (Table 5, Supplementary Material). The economic value of the ecosystem service ‘climate regulation’ was calculated by multiplication of the mean net carbon flux in the soil per hectare and year (total organic soil carbon mass at year x – total organic soil carbon mass at the beginning of the experiment in year 2013) * 3.66 (transformation factor from carbon mass to CO_2_ mass based on the atomic mass of carbon and oxygen) with an accounting price of 195 € / t of CO_2_. This accounting price is recommended by the German Federal Environmental Agency for assessing environmental costs (German Federal Environmental Agency, 2021), and is meant to capture the social cost of carbon, i.e., the aggregate damages of emitting a ton of carbon dioxide into the atmosphere. Transformation of the percentage specification of total organic carbon in the soil mass into carbon mass per area was done based on the assumption of a carbon distribution in alifsols up to a depth of 30 cm (Sulman et al., 2020) and a soil weight of 1350 kg/m^3^ (Engineering ToolBox, 2008).

The economic value of the ecosystem service ‘water regulation’ was calculated by multiplication of the annual net nitrogen flux (kg nitrogen surplus / ha) with an accounting price capturing the social cost of excess nitrogen in agroecosystems. Due to the geological site conditions, especially the occurrence of slack water (surface water accumulated on an impermeable or less permeable soil layer) (Geodatenportal Sachsen-Anhalt, 2022), a social cost value of 7.30 € / kg N_r_ surplus is used as the social cost value for N_r_ leaching into surface water bodies, again following the guidelines of the German Federal Environmental Agency for assessing environmental costs (German Federal Environmental Agency, 2021).

The insurance value of the ecosystem services ‘biodiversity conservation’ and ‘soil health’ was calculated according to Baumgärtner (2007). It is based on evidence obtained from our experimental data that both biodiversity (R^2^ = 0.197, p<0.01**) and soil health (R^2^ = 0.272, p<0.001***) correlate with yield stability which is an underlying assumption so that an insurance value can be calculated. The risk premium *RP* of the ecosystem function ‘yield’ is calculated on plot level as the mean value of the yield 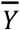 minus the certainty equivalent, with yield coefficient of variation *CV* and risk aversion *r*, assuming that the yield is lognormally distributed. A value of 1.26 was chosen for the relative risk aversion *r* as the median of 58 studies assessing relative risk aversion according to Elminejad et al. (2022).

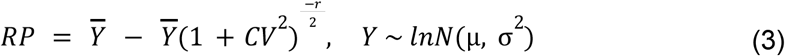

Risk premium was shown as a function of biodiversity *b* and soil health *h*. A multivariate, polynomial regression model of the form of RP(*b,h*) = x_1_ + x_2_ *b* + x_3_ *b*^2^ + x_4_ *h* + x_5_ *h*^2^ + x_6_ *b h* was used to calculate the response surface (Figure 8, Supplementary material). Then, the insurance value *I*_*B*_ of biodiversity at biodiversity level *b* and soil health level *h* is given as the difference between risk premium *RP* at biodiversity level 0 and soil health level *h* and the risk premium *RP* at biodiversity level *b* and soil health level *h*:

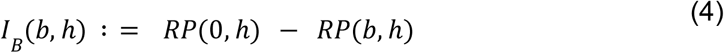

**Figure 8.**
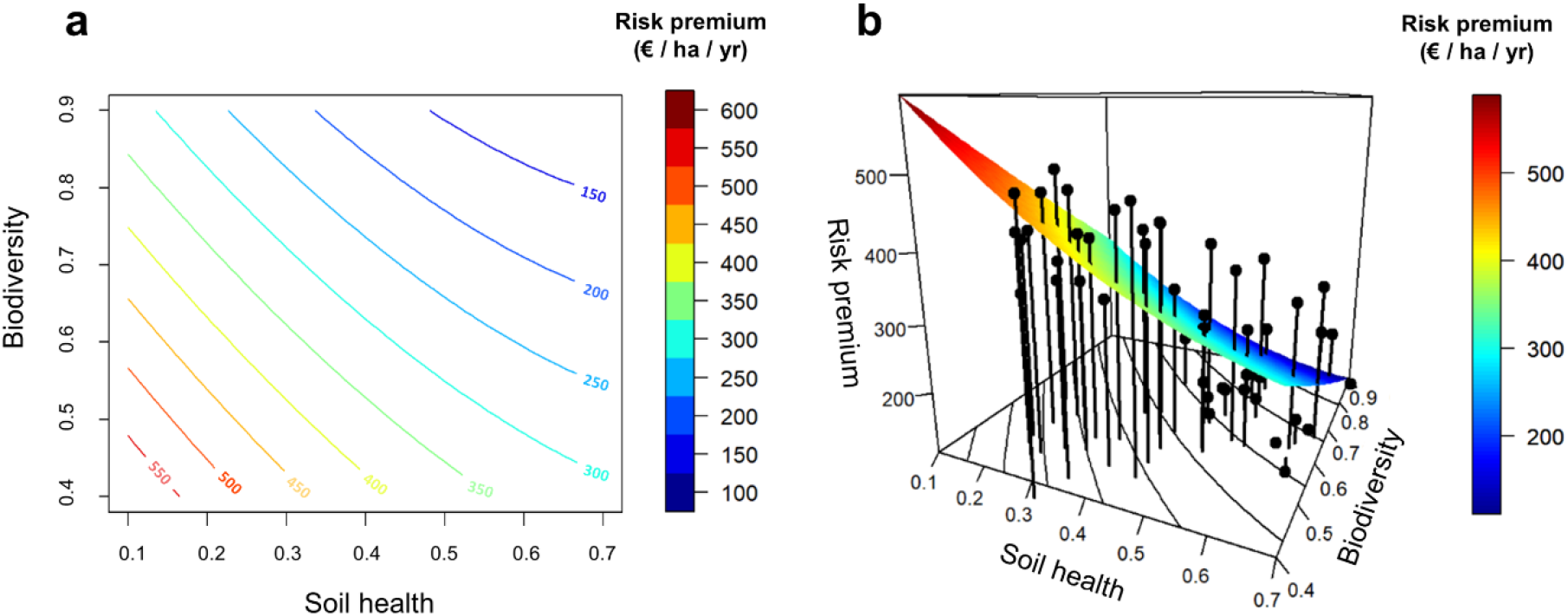
Risk premium as function of biodiversity and soil health (normalised levels) for a risk aversion value of r = 1.26 as median of 58 studies assessing relative risk aversion (Elminejad et al., 2022): response surface used for the calculation of the insurance value of biodiversity and soil health in two (a) and three dimensions (b).

**Figure 9.**
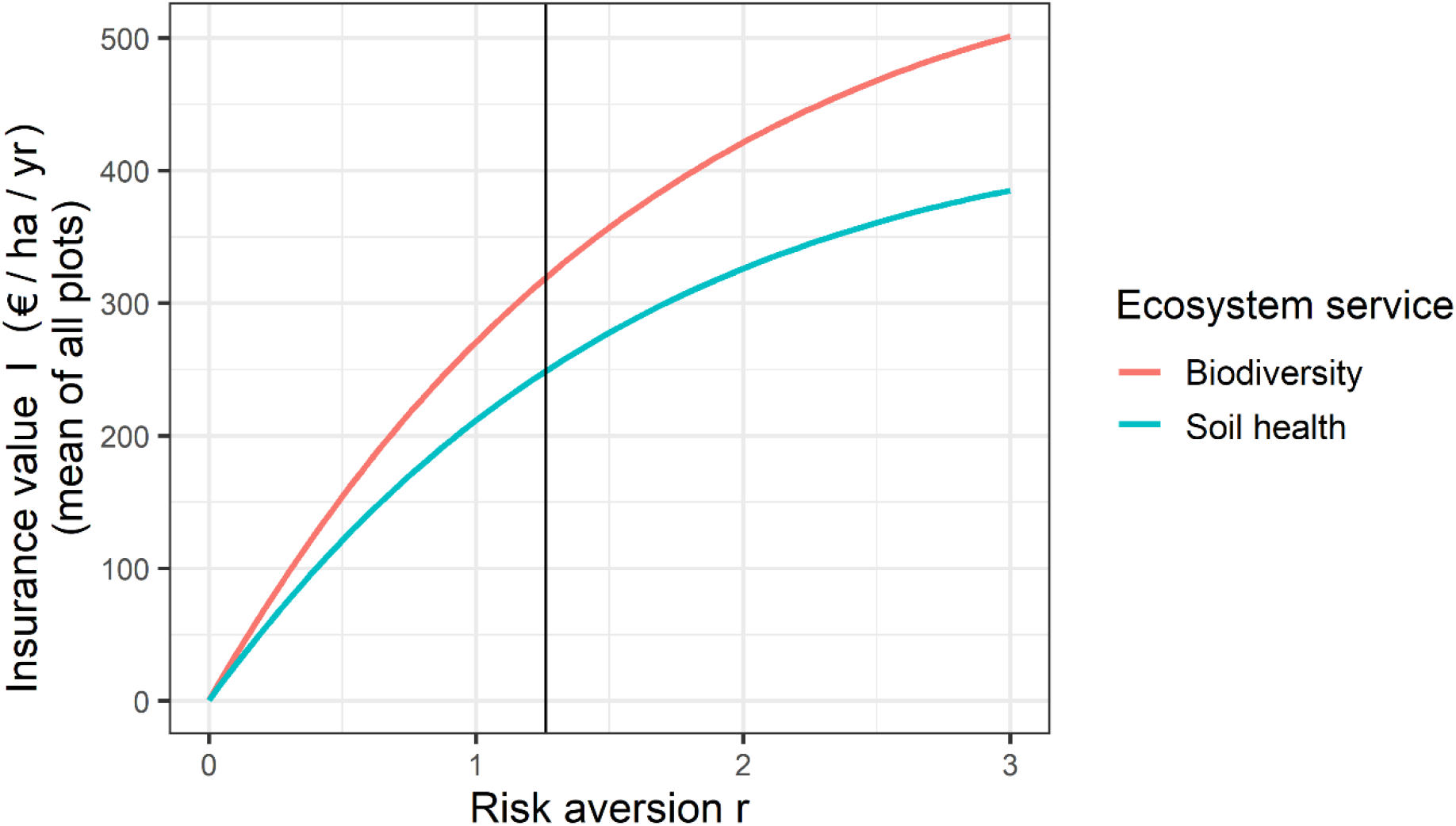
Insurance value (mean of all plots) as function of risk aversion for both biodiversity and soil health. Vertical bar indicates the value of risk aversion r that was chosen for this study (1.26 as median of 58 studies assessing relative risk aversion (Elminejad et al., 2022)).

Vice versa, the insurance value *I*_*H*_ of soil health at biodiversity level *b* and soil health level *h* is given as the difference between risk premium *RP* at biodiversity level *b* and soil health level 0 and the risk premium *RP* at biodiversity level *b* and soil health level *h*:

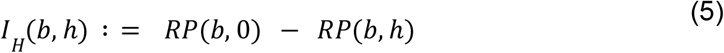

so that the total insurance value *I*_*HB*_ of the two insurance providing ecosystem services is given as the difference between risk premium *RP* at biodiversity level 0 and soil health level 0 and the risk premium *RP* at biodiversity level *b* and soil health level *h*:

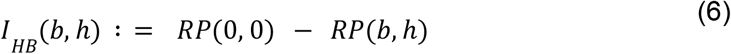

The economic ecosystem multifunctionality value *eEMF* was calculated by summing the values of the single ecosystem services, which are obtained by multiplication of the level of the respective ecosystem service *ES*_*i*_ with the accounting price *p*_*i*_ of the respective ecosystem service.

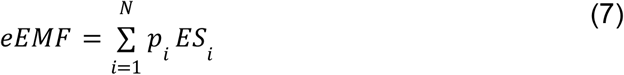

### 5.8. Statistical analysis

All calculations and the statistical analysis were conducted using R (R Core Team, 2021) with the packages ‘extrafont’, ‘ggplot2’, ‘ggpubr’, ‘lme4’, ‘lmerTest’, ‘car’, ‘multcomp’, ‘multcompView’, ‘stringr’, ‘dplyr’ and ‘plot3D’. See supplementary material for the complete R code and the dataset that was compiled for the analysis. Statistical analysis of the level of ecosystem functions and EFM was conducted using Type III ANOVA Satterthwaite’
ss method in accordance with the split-plot design of the experiment.

## 6. Additional information

### Supplementary material

comprises information regarding individual ecosystem function responses to different climate and land-use type types, additional figures and tables as well as a document describing the realisation of all calculations with R and a document explaining the abbreviations used within the calculations.

## Acknowledgement

We thank the staff of the Bad Lauchstädt Experimental Research Station and the GCEF (especially Konrad Kirsch) for the maintenance of the study area and Stefan Klotz and François Buscot for their role in setting up the GCEF. We thank Elke Schulz for the provision of data on microbial biomass, plant available nitrogen and total organic soil carbon. We thank Gabriele Rada for supporting the data visualisation.

## Author Contributions

F.S., N.E. & M.Q. conceptualised and designed the research; M.S. & HA designed the GCEF; M.S. is the scientific coordinator of the GCEF; M.S., T.R., R.Y., H.A., I.M., S.B., E.B., C.R., S.H., J.S. & M.C. collected the data; F.S. synthesised the data; F.S. analysed the data; F.S. wrote the first draft of the manuscript; F.S., N.E. & M.Q. were the core writing team; all co-authors contributed to revisions of the manuscript.

## 8. Supplementary material

### 8.1 Ecosystem function responses to different climate and land-use types

Yield is negatively affected by future climate and, for grassland, significantly higher under intensive management than under sustainable management. Surprisingly, for farmland, yield is higher under sustainable management than under conventional (i.e., intensive) farming. Total organic soil carbon is lower for farmland than for grassland. Nitrogen surplus is significantly higher for the intensively-managed land-use types and in general increased under future climate. Microbial biomass is lower for farmland and shows an interaction effect: future climate increases microbial biomass only in extensive pasture (EP). Effects of land-use type and future climate differ across individual enzymes: cellulase activity is decreased under future climate, whereas no land-use effect is found. N-acetylglucosaminidase activity is lower for farmland compared to grassland, and not significantly affected by future climate. Acid-phosphatase shows a lower activity only for sustainably-managed farmland. Belowground decomposition rate for grasslands is significantly lower under intensive management, while for farmland no difference was found between the management types. Aboveground (microbial) decomposition rate is lower in farmlands than in grasslands. Aboveground (microbial + faunal) decomposition rate is neither affected by climate nor land-use type and intensity. Soil mesofauna diversity is not significantly affected by any treatment, whereas soil macrofauna diversity is lower for farmland compared to grassland. Soil nematode diversity is significantly decreased in intensively-managed grassland compared to all other land-use types.

